# Kombucha-Derived Cellulose Non-wovens: Growth Optimization, Mechanics, and Recycling

**DOI:** 10.64898/2026.06.09.730694

**Authors:** Luying He, Julia E. Lopez, Jessica D. Schiffman

**Author notes:** Corresponding author: Jessica D. Schiffman.

## Abstract

The environmental impact of synthetic textiles has prompted the search for sustainable and biodegradable alternatives. This study correlates the growth conditions used to produce kombucha-derived cellulose non-woven mats with their mechanical performance as a function of post-processing. Systematically, the fermentation and growth parameters of the non-wovens, including inoculum density, carbon-source loading, temperature, and pH value were investigated. Thick, uniform non-wovens were obtained using mildly acidic conditions that balanced nutrient availability and growth rate, moderate inoculum and carbon loading at 30 °C. Next, we used uniaxial tensile testing and rheology to thoroughly compare the mechanical properties of two post-processing routes, lyophilization and oven-drying against the as-produced wet non-wovens. Overall, the lyophilized non-wovens displayed the highest ultimate tensile strength (14.36 ±0.9 MPa) and elongation at break (24.54 ±1.9%), which were statistically greater than the oven-dried (2.54 ±0.3 MPa, 6.03 ±0.8%) and the wet non-wovens (1.66 ±0.3 MPa, 9.35 ±2.8%). We conclude by performing a proof-of-concept recyclability experiment: we showed that kombucha-derived clothing could be enzymatically degraded and then re-manufactured into new nanofibers by electrospinning. Together, these results demonstrate a circular pathway encompassing the growth and processing of mechanically robust kombucha-derived cellulose non-wovens, as well as their biodegradation and re-manufacturing.

## INTRODUCTION

Globally, textile production generates ∼92 million tons of waste each year, yet less than 1% is recycled into new clothing.^1^ In 2023, synthetic polymers such as polyester, nylon and acrylic accounted for 60% of the global textile production^2,3^ and their manufacturing used 50-100 million tons of crude oil, *i.e.*, ∼1% of the annual global oil usage. Fiber production is also highly carbon-intensive, with polyester production releasing around 6 kg CO₂ per t-shirt, which is approximately triple that of cotton.^4^ Most synthetic clothing does not easily degrade and thus, it persists in the environment for decades.^5^ These challenges underscore the need for sustainable, circular textile platforms.

Researchers have turned to natural polymers such as cellulose, including bacterial cellulose, as sustainable alternatives,^6^ however, sourcing sufficient feedstock, large-scale production, and high costs remain significant challenges. Microbial production of bacterial cellulose is particularly resource-intensive, requiring nutrient-rich media and controlled bioprocessing, which keeps it far more expensive than plant-derived cellulose.^7,8^ Static cultures yield coherent sheets but they are only ∼1 wt% solid and labor-intensive to scale, whereas submerged fermentation is more scalable yet often produces pelletized aggregates and cellulose-deficient mutants that depress yield. In both modes, cellulose nanofiber diameter, membrane thickness, and crystallinity are highly sensitive to strain and culture conditions, complicating quality control.^9–12^

Kombucha-derived cellulose non-wovens are complex fiber networks produced from a microbial consortium composed of yeasts and multiple bacterial strains.^13^ More specifically, during the fermentation of kombucha tea, yeasts cleave sucrose into glucose and fructose, which Acetobacter, typically *Komagataeibacter xylinus,* use to generate acetic acid and polymerize cellulose microfibrils.^14^ Over a 14 day period, a mat of bacterial cellulose is produced and floats at the air-liquid interface.^13,15,16^ Previously, researchers have demonstrated that the type of tea (*e.g.*, black, green, herbal) and fermentation conditions (*e.g.,* sugar concentration, temperature, time) steer metabolic flux distribution and extracellular polymer deposition, thereby controlling the fiber color, antioxidant capacity, and growth kinetics.^14^ Alternative carbon substrates, including agro-industrial byproducts, nitrogen sources, and co-culture strategies can also shift osmotic balance, productivity, and mechanical robustness through cross-feeding and controlled acidification, again translating upstream metabolic changes into downstream non-wovens structure.^13,17^ Although kombucha-based non-wovens have been explored, most of the reports have focused on how the starting ingredients and substrates influence the production of kombucha tea, cellulose production, and biomedical applications.^18–20^ A systematic optimization of the most relevant fermentation variables, and a thorough materials characterization of the optimized, post-treated non-wovens remain largely unexplored, as does the fit of these materials into a circular economy.

The goal of this work is to first provide a universal recipe that produces thick kombucha-derived cellulose non-wovens from a readily available commercial starter kit comprised of symbiotic culture of bacteria and yeast (SCOBY), as illustrated in **Figure 1**. The impact of how classic fermentation parameters, including inoculum ratio, sucrose and black tea concentration, temperature, and pH value influence non-woven thickness and yeast cell density over 14 days was investigated. Next, we continued to produce cellulose non-wovens using the optimized conditions and explored the impact of non-woven thickness, as well as post-processing via oven-drying and lyophilization on their mechanical properties using both uniaxial tensile testing and rheological characterization. Our findings were benchmarked against other mechanical properties reported in the literature. Finally, as a proof-of-concept demonstration, we first fermented kombucha-derived cellulose into the shape of baby clothing; next, we used enzymatic degradation to break down the textiles into a new precursor solution, which we re-manufactured into nanofiber scaffolds. In this work, we aim to demonstrate the potential of kombucha-derived cellulose non-wovens as a robust platform that fits within a circular textile economy for applications including clothing, separation membranes, green packaging, and wound dressings.

**Figure 1.**
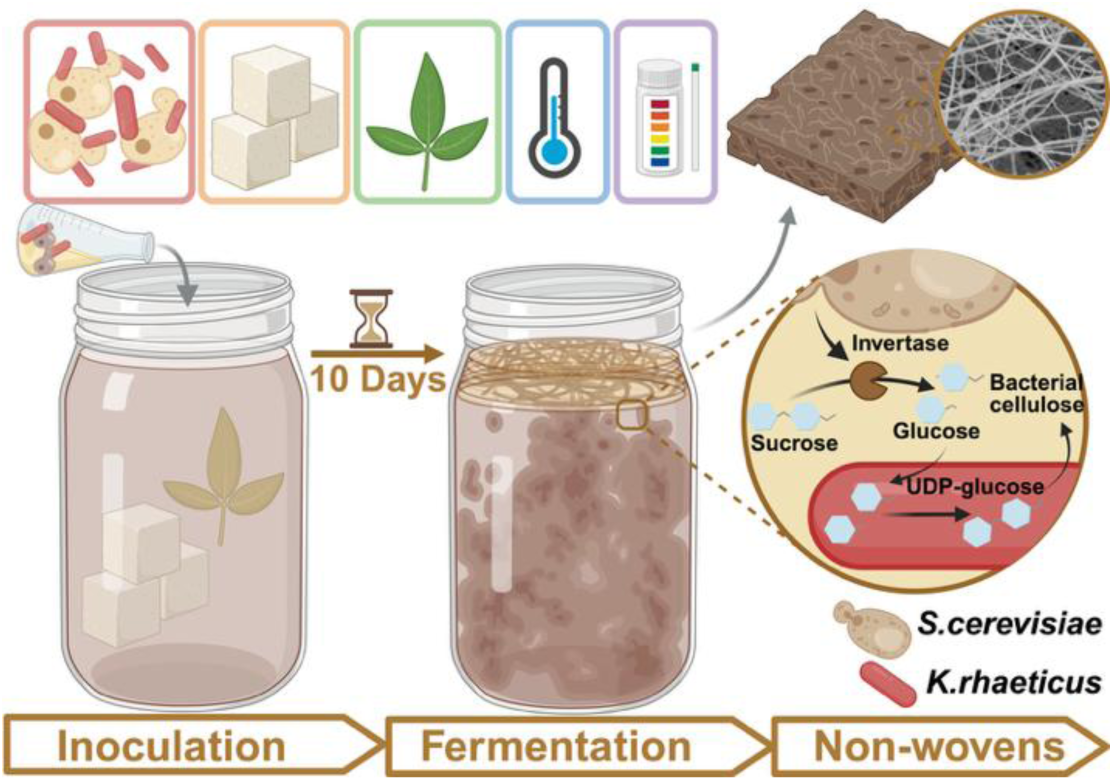
Schematic diagram of the process used to grow kombucha-derived cellulose non-wovens. The specific parameters that were tested are shown pictorially and include: inoculum, sucrose and tea concentration, temperature, and pH value. Details are provided in **Table 1**.

**Table 1.**
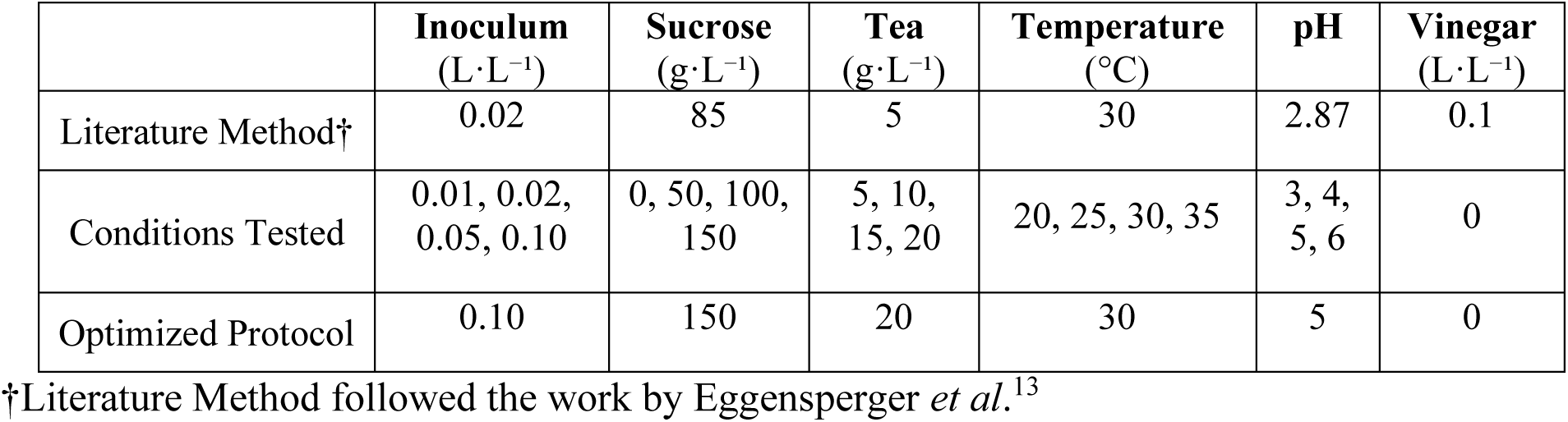
Kombucha Fermentation Parameters Evaluated in This Work.

## MATERIALS AND METHODS

### Materials and Chemicals

All chemicals were used as received without further purification. Black tea leaves (Camellia sinensis) and SCOBY were purchased from The Kombucha Shop (Madison, WI). White vinegar was purchased from Walmart (Bentonville, AR). Ethanol (200 proof) and sodium chloride (NaCl) were purchased from Fisher Scientific (Hampton, NH). Sucrose (≥99.0%, ACS reagent, Cat. No. S7903), yeast extract (bacteriological grade, Cat. No. Y1625), peptone (bacteriological grade, Cat. No. P7000), D-(+)-glucose monohydrate (dextrose, ≥ 99.0%, Cat. No. G8270), agar (bacteriological grade, Cat. No. A1296), polyvinyl alcohol (PVA, ≥99% hydrolyzed, Cat. No.563900) were purchased from Sigma-Aldrich (St. Louis, MO, USA). Phosphate-buffered saline (PBS) was prepared using NaCl (≥99.0%), potassium chloride (KCl, ≥99.0%), disodium hydrogen phosphate (Na₂HPO₄, ≥99.0%), and potassium dihydrogen phosphate (KH₂PO₄, ≥99.0%) purchased from Fisher Scientific (Hampton, NH, USA). Cellulase from *Trichoderma reesei* (≥ 1000 U g^-1^; Cat. C2730), sodium acetate anhydrous (NaOAc, ≥ 99%; Cat. S2889), glacial acetic acid (AcOH, ≥ 99.7%; Cat. 695092), and calcofluor white stain (CFW; cellufluor, ≥ 90%; Cat. 18909) were purchased from Millipore-Sigma (St. Louis, MO, USA). Deionized (DI) water (18.2 MΩ·cm) was produced using a Barnstead Nanopure Infinity water purification system (Thermo Fisher Scientific, Waltham, MA, USA).

### Fermentation of Kombucha-Derived Non-Wovens

As displayed on **Table 1**, a systematic study was conducted to determine the optimal conditions to grow non-wovens. The parameters that were varied included the kombucha inoculum (0.01-0.1 L·L⁻¹), sucrose (0-150 g·L⁻¹), tea concentration (5-20 g·L⁻¹), temperature (20-35±0.5°C), and the initial pH value (3-6). The control “literature” method, followed the general approach of Eggensperger *et al.*^13^ wherein the fermentation medium (1 L total volume) contained 85 g·L⁻¹sucrose and 0.1 L·L⁻¹ distilled white vinegar (pH = 2.87) prepared in black tea infusion (4.6g tea in 900mL boiling DI water, steeped 1 h, cooled to 30°C, filtered). The jars were inoculated with 0.02 L·L⁻¹ kombucha inoculum (10⁷ CFU mL^-1^ total acetic acid bacteria and yeast) without a SCOBY and covered with two layers of autoclaved gauze to permit oxygen diffusion.^13^ All media were prepared using DI water that was sterilized by autoclaving (121°C, 20 min). Cultures were maintained statically for 14 days with three independent replicates per condition. Non-wovens were retrieved using flame-sterilized tweezers, rinsed three times with DI water, and blotted to remove extra liquid using sterile gauze before being stored in DI water at 4°C in DI water. Samples were used within 48 h. All glassware and utensils were steam-sterilized, and negative (uninoculated) controls confirmed asepsis.

### Enumeration of Saccharomyces cerevisiae

Yeast Peptone Dextrose (YPD) agar plates were prepared by suspending 10 g yeast extract, 20 g proteose peptone, 20 g dextrose, and 15 g agar in 1.0 L of DI water. The slurry was heated to a gentle boil with constant stirring until all solids dissolved. The pH value was adjusted to 6.0 ±0.1 with sterile 1 M HCl or 5 M NaOH, and the medium was sterilized by autoclaving at 121°C for 20 min (15 psi). After cooling to 55°C, ∼25 mL of molten agar was aseptically dispensed into sterile 90 mm Petri dishes inside a biosafety cabinet, allowed to solidify, inverted, and stored at 4°C for ≤7 days. PBS was prepared by dissolving 8.0 g NaCl (≥99.0%), 0.2 g KCl (≥99.0%), 1.44 g Na₂HPO₄ (≥99.0%), and 0.24 g KH₂PO₄ (≥99.0%) in 1 L of DI water with magnetic stirring at room temperature. The pH value was adjusted to 7.40 ±0.05 using 1 M HCl or NaOH, and the solution was sterilized by autoclaving at 121°C for 20 min.

For CFU determination, yeast cultures were sampled at the desired time points, vortex-mixed, and serially diluted (10× series) in sterile 1× PBS (pH 7.4). Aliquots (100 µL) of the appropriate dilutions were spread in triplicate onto YPD agar plates and oven-dried. Plates were incubated at 30°C for 24-48 hr, after which colonies in the 30-300 range were counted. CFU values were calculated as **Equation 1**:

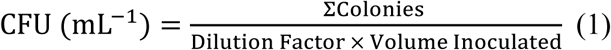

Biological duplicates and medium-only negative controls were included in every assay to verify sterility and reproducibility.

### Post-Processing and Characterization of Kombucha Derived Non-Wovens

The as-fermented non-wovens were collected and evaluated (1) wet (*i.e.*, without further processing), (2) oven dried at 98°C for 24 hr (using a Thermmax Scientific Products, Model CCR-2-C-0-R1-0-DX-EL-UL Oven, Colmar, PA, USA), or (3) lyophilization (Labconco FreeZone 2.5 Plus) after being placed in a −80°C freezer for 12 hr. A drying temperature of 98 °C was chosen because it is close to, but just below the boiling point of water. The thickness of the non-wovens was measured at five random positions with a Mitutoyo 293-330 digital micrometer (resolution = 1 µm; Toronto, Ontario, Canada). Normalized thickness was determined using **Equation 2**.^21^

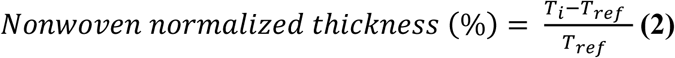

where *T_i_* is the thickness of each sample and *T_ref_* is the thickness of the reference condition within each culture-variable group. Thickness values are reported as mean ±standard deviation (n = 15). Fibers within the non-wovens were imaged using scanning electron microscopy (SEM, FEI Magellan 400 XHR-SEM) at 5 kV at a working distance of 4 mm. Before imaging, samples were mounted on carbon adhesive tape and sputter-coated with a 2 nm gold layer (Cressington 208HR) to enhance the electron conductivity. Using FIJI (ImageJ, v1.53a),^22^ ≥50 random fiber diameters were measured across multiple fields of view. A two-tailed, unpaired t-test (Orign.Ink) was used to assess differences among groups (p<0.05 considered significant). A Bruker Alpha (Bruker Corporation, Billerica, MA) instrument was used to conduct attenuated total reflectance Fourier transform infrared (ATR-FTIR) of kombucha derived non-wovens and commercial cellulose powder controls. Spectra were collected at a resolution of 4 cm^-1^, averaged over 24 scans over a range of 4000-400 cm^-1^.

### Mechanical Testing of Kombucha-Derived Non-Wovens

Rheological measurements were performed following established protocols.^23^ Tests were conducted on a Kinexus Pro rheometer (Malvern Instruments) with a parallel-plate geometry (20 mm diameter, 0.5 cm gap) at 25°C. Samples used for rheology were 25.4 mm in diameter and at least 0.2 mm thick. Small-amplitude oscillatory shear (SAOS) sweeps were used to measure the storage modulus (G’) and loss modulus (G”) from amplitudes. Frequency sweeps were carried out within the linear viscoelastic regime to ensure that the reported moduli were independent of strain amplitude. More details about the oscillatory rheological method are provided in the **Supporting Information**.

Uniaxial tensile tests were performed using a texture analyzer (TA.XTPlus, Stable Micro Systems Ltd., Hamilton, MA) equipped with mechanical grips at a crosshead speed of 1.0 mm/s. Non-woven samples were cut into dog-bone samples (gauge length=25 ±0.47 mm, width=4.8 ±0.076 mm) in accordance with ASTM D882 (ISO 527-3)^24^ using a laser cutter (VLS 3.50 30W CO_2_ laser, Universal Laser Systems Inc.) using 70% power, 2.5% speed, and 1000 PPI. The laser-cutting did not char, darken, or thermally deform the samples. Sample thickness and width were measured at five locations using a Mitutoyo 293-330 digital micrometer. More details about how the Young’s modulus, ultimate tensile strength (UTS), elongation at break, strain energy density to fracture (W), and fracture surface energy (G) were determined and calculated are provided in the **Supporting Information**, including **Equations S1** and **S2**. At least four independent replicates were tested for each condition. Statistical comparisons between drying treatments were performed using two-tailed Student’s t-tests.

### Biodegradation of Kombucha-Derived Non-Wovens

Wet non-wovens were punched into circular specimens (Ø=25 mm, 2 mm thick) using a stainless-steel biopsy punch. A 50 mM sodium acetate/acetic acid buffer in DI water (pH 4.80 ±0.05) was prepared and sterilized using a 0.22 µm PES membrane filter.^25^ Cellulase was dissolved in the same buffer to prepare a 10 U mL⁻¹ stock solution and diluted to a working concentration of 6 U mL⁻¹ before use.^26^ Non-wovens were immersed in either the buffer alone (control) or cellulase solution (6 U mL⁻¹ in the same buffer) and incubated for up to 72 hr. After degradation, the remaining solids (50 mL) were collected by vacuum filtration and lyophilized prior to re-processing. Digital images were collected to monitor degradation.

### Re-manufacturing via Electrospinning

The recovered lyophilized solids were mixed with 2 mL of an aqueous 10% (w/v) PVA solution (equivalent to 0.20 g PVA) and cellulose stain (0.1 mg mL⁻¹ CFW) by continuous stirring at room temperature for 12 h. The mixture was loaded into a pre-sterilized disposable 5 mL syringe (Fisher Scientific, Hampton, NH) for electrospinning using our in-house system. Briefly, an aluminum foil wrapped copper sheet (15×15×0.32 cm) was placed 18 cm from the needle tip and used as the collector plate. The applied voltage used was 17.5 kV (Gamma High Voltage Research, Ormond, FL), flow rate of 0.50 mL h⁻¹ was used.^27^ Environmental conditions were controlled at 23°C and 22% relative humidity using an environmental chamber. Images of the electrospun samples were acquired using ZEN 2.3 Pro software and a Zeiss Axio Imager A2m fluorescence microscope equipped with a blue fluorescence filter set (excitation 350-400 nm; emission 420-500 nm).

### Statistical Analysis

Data are presented as mean ±standard deviation. For two-group comparisons, an unpaired, two-tailed Student’s t-test was used. For multi-group comparisons (*e.g.*, different pH value or sugar concentrations), one-way ANOVA followed by Tukey’s post-hoc test was conducted (α = 0.05). All statistical analyses were performed using OriginPro 2024 or Microsoft Excel 2024.

## RESULTS AND DISCUSSION

### Influence of Culture Variables on Kombucha Derived Non-Wovens Thickness

We first examined how a library of different starting fermentation parameters impacted the development of kombucha-derived non-wovens over a 14-day growth period. A key feature of our microbial consortium is that the cellulose-producing bacteria are unable to directly metabolize sucrose and instead depend on yeast-mediated hydrolysis to generate fermentable monosaccharides.^28,29^ As yeast populations grow, extracellular enzymatic activity converts sucrose into glucose, thereby supplying the primary carbon source required for bacterial cellulose biosynthesis.^30,31^ Increasing sucrose concentration effectively increases carbon availability to the microbial consortium, enhancing metabolic flux toward cellulose formation and accelerating the growth of non-wovens (**Figure 2A-B**).^32^ This interdependence means that altering parameters, such as inoculum, sugar concentration, or nutrient composition tunes fermentation kinetics and, in turn, the rate of cellulose accumulation at the air-water interface.

**Figure 2.**
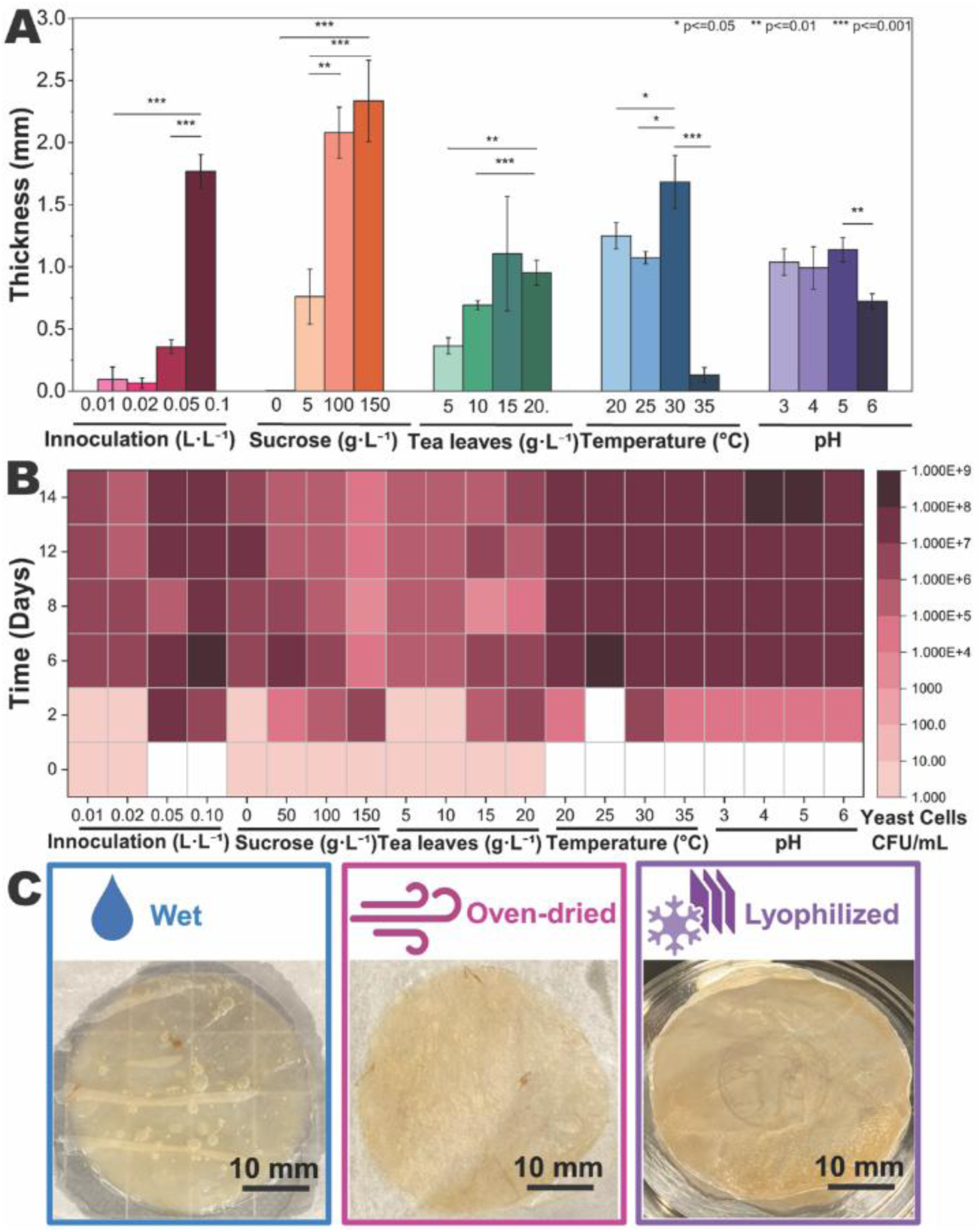
Impact of culture variables, including inoculum, sucrose, tea leaf, temperature, and pH value on **(A)** kombucha derived non-wovens thickness and **(B)** yeast cell density. The white boxes indicate conditions under which yeast cell density was below the assay’s limit of detection. (**A** and **B**) The fermentation period was 14 days. **(C)** Representative digital images of kombucha-derived cellulose non-wovens: as-prepared (wet), oven-dried, and lyophilized. A 10 mm scale bar is provided.

As shown on **Figure 2A** and tabulated on **Table S1**, it was observed that when the inoculum concentration was increased from 0.01 L·L⁻¹ to 0.10 L·L⁻¹ there was an ∼18× increase in non-woven thickness, from 0.096 ±0.08 mm to 1.769 ±0.11 mm. At lower inoculate levels (0.01 to 0.05 L·L⁻¹), limited yeast abundance slows invertase-mediated sucrose hydrolysis and prolongs the microbial lag phase, constraining early metabolic flux into cellulose synthesis.^33,34^ In contrast, a 0.10 L·L⁻¹ inoculum provides sufficient active biomass to rapidly convert sucrose into fermentable sugars, shorten the lag phase, and sustain high cellulose production rates, resulting in statistically thicker non-wovens. When the higher inoculum was used, the resultant increase in non-woven thickness suggests that there was a threshold effect in consortium activity rather than a linear biomass scaling. A positive correlation emerged between sucrose levels and thickness: the greatest sucrose concentration yielded the thickest non-wovens, which was consistent with a previous report that indicated that higher sugar availability can accelerate bacterial and yeast proliferation.^37^ In this study, we did not explore even higher sucrose concentrations to avoid imposing osmotic or inhibitory stress on the microbial consortium and to maintain relevance to scalable fermentation conditions. Prior studies have reported that excessive sugar levels can suppress microbial activity, alter community balance, and reduce cellulose productivity despite increased carbon availability.^36^

Raising the tea leaf concentration from 5 g·L⁻¹ to 20 g·L⁻¹ yielded a 26% increase in thickness, from 0.364 ±0.05 mm to 0.952 ±0.08 mm, suggesting that increasing the tea concentration provides additional growth-supporting nutrients and bioactive compounds that enhance consortium activity and cellulose formation. This interpretation is consistent with prior studies showing that tea-derived nutrient composition and carbon sources affect bacterial cellulose growth. Yim *et al*.^37^ reported that varying tea type and sugar source altered non-wovens yield and thickness, with the greatest thickness (0.213 mm) obtained using green tea and sucrose.^37^ Notably, using very high tea concentrations can introduce excess phenolics or insoluble residues that inhibit microbial activity and complicate downstream processing.

To relate membrane growth to consortium activity, **Figure 2B** shows a heatmap of yeast cell population under the same culture variables examined in **Figure 2A** over a 14-day incubation period. We note that under some conditions during days 0-1 the yeast cell densities were below the assay detection limit and therefore are displayed as a white box. From the yeast CFU heatmap in **Figure 2B**, the 50 g L⁻¹ sucrose condition exhibits a canonical microbial growth trajectory over 14 days. Yeast populations increase from 10⁴-10⁵ CFU mL⁻¹ on days 0-2 to a peak of 10⁷-10⁸ CFU mL⁻¹ on days 2-6, followed by a progressive decline to 10⁶-10⁷ CFU mL⁻¹ on days 6-8 and 10⁵-10⁶ CFU mL⁻¹ by day 14, indicating a transition from exponential growth to post-peak decay (**Figure 2B**). Yeast cell densities at day 14 were substantially higher in cultures supplemented with 20 g·L⁻¹ tea leaves (10⁶-10⁷ CFU·mL⁻¹) than those containing 5 g·L⁻¹ tea leaves (10⁵-10⁶ CFU·mL⁻¹). This enhancement in yeast growth is attributed to the increased availability of tea-derived micronutrients and polyphenols, which are known to support yeast metabolic activity and stress tolerance during fermentation.^42,43^ In this study, yeast density was monitored as an accessible indicator of fermentation activity and community stability, while non-woven thickness served as the primary material-level output of cellulose accumulation. Acetic acid bacteria are the primary cellulose-producing microorganisms in kombucha cultures, and we are not interpreting yeast density as a direct proxy for cellulose-producing bacterial abundance. Direct quantification of acetic acid bacteria by selective plating, qPCR, or sequencing-based approaches will be valuable in future work to further connect microbial community structure with cellulose non-woven formation.

We next examined how environmental stress modulates cellulose synthase activity and consortium dynamics. Non-wovens growth was maximized at 30°C, whereas at 35°C, cellulose accumulation was markedly reduced despite sustained yeast populations (**Figure 2A-B**). The thickest non-woven, 1.683 ±0.17 mm was produced using an incubation temperature of 30°C, while only a 0.130 ±0.05 mm non-woven developed at 35°C, possibly due to heat stress on the microbial consortia (**Figure 2A**). The yeast cell density was equivalent across the 20-35 °C temperature range. This is consistent with prior reports demonstrating that the cellulose synthase complex of *K. rhaeticus* rapidly loses activity above 35°C due to thermal aggregation of the CesA subunit when not stabilized by the cytoplasmic membrane, thereby limiting efficient cellulose biosynthesis at elevated temperatures.^40,41^

Fermentation pH strongly modulated fiber mat formation and culture stability. Mildly acidic conditions with a pH value of 4 or 5 promoted robust cellulose accumulation and suppressed microbial contamination, whereas increasing the initial pH value to 6 significantly hindered non-woven growth (**Figure 2A-B**). This behavior reflects the acid tolerance and optimal activity of cellulose-producing acetic acid bacteria where acidic conditions remain compatible with yeast metabolism, enabling effective metabolic partitioning within the consortium and sustained cellulose accumulation.^42^ A mildly acidic pH value of 5 produced significantly thicker non-wovens measuring 1.145 ±0.11 mm, versus a pH value of 6 (0.722 ±0.06 mm).

Non-wovens thickness reflects both the initial fermentation conditions, as well as the temporal dynamics and spatial organization of the consortium. In co-culture, yeast and *K. rhaeticus* exhibited comparable cell density (10⁶-10⁷ CFU·mL⁻¹), while *K. rhaeticus* was preferentially enriched within the cellulose non-wovens, reflecting coupled yet spatially partitioned growth.^43^ Non-wovens thickening continued even after the yeast count reached a steady state. This sustained cellulose accumulation is attributed to prior sucrose hydrolysis by yeast, including cells embedded within the developing fibers, which generates sufficient glucose to support continued bacterial cellulose synthesis despite declining planktonic yeast abundance.^44^

### Characteristics of Kombucha-Derived Non-Wovens

Next, we produced consistent non-wovens using the optimized protocol (described in **Table 1**) to explore the role that post-processing has on their morphology and mechanical properties. As displayed in the digital images provided in **Figure 2C**, the kombucha-derived membranes were (i) used as-prepared (wet), or post-treated via (ii) oven-drying at 98 °C, or (iii) lyophilization. The as-prepared, *i.e.*, non-treated non-woven appeared to be a swollen translucent gel. In contrast, the oven-dried non-wovens appear more flattened and more transparent, whereas the lyophilized samples were opaque and white. Indeed, Sozcu *et al.*^45^ observed that oven-dried bacterial cellulose films become transparent with collapsed fibers, whereas freeze-dried films stay opaque and porous.^45^

SEM micrographs provided in **Figure 3** display the morphology of the kombucha-derived non-wovens. Because SEM requires a dry sample, our visualization focused on the lyophilized samples. We observed porous mats composed of ultrafine fiber bundles interwoven in random orientations. The highly porous, non-woven fiber architecture provides high surface area. Individual fibers had an average diameter of 0.06 ±0.04 *μ*m (**Figure 3E**), and the range of fiber diameters observed were between 0.02 *μ*m to 0.18 *μ*m. Bacterial-produced cellulose nanofibers have previously been reported to have diameters ranging from 0.02-0.10 *μ*m.^16,46–48^ Our fibers were on par with those previously reported, differences in fiber diameters can result from bundling or co-aggregation due to the multi-strain environment in kombucha fermentation.^49^ The fiber diameters distribution shown in **Figure 3E** suggests that we had a homogeneous biosynthesis process because large microfibrils or bundles were not observed.

**Figure 3.**
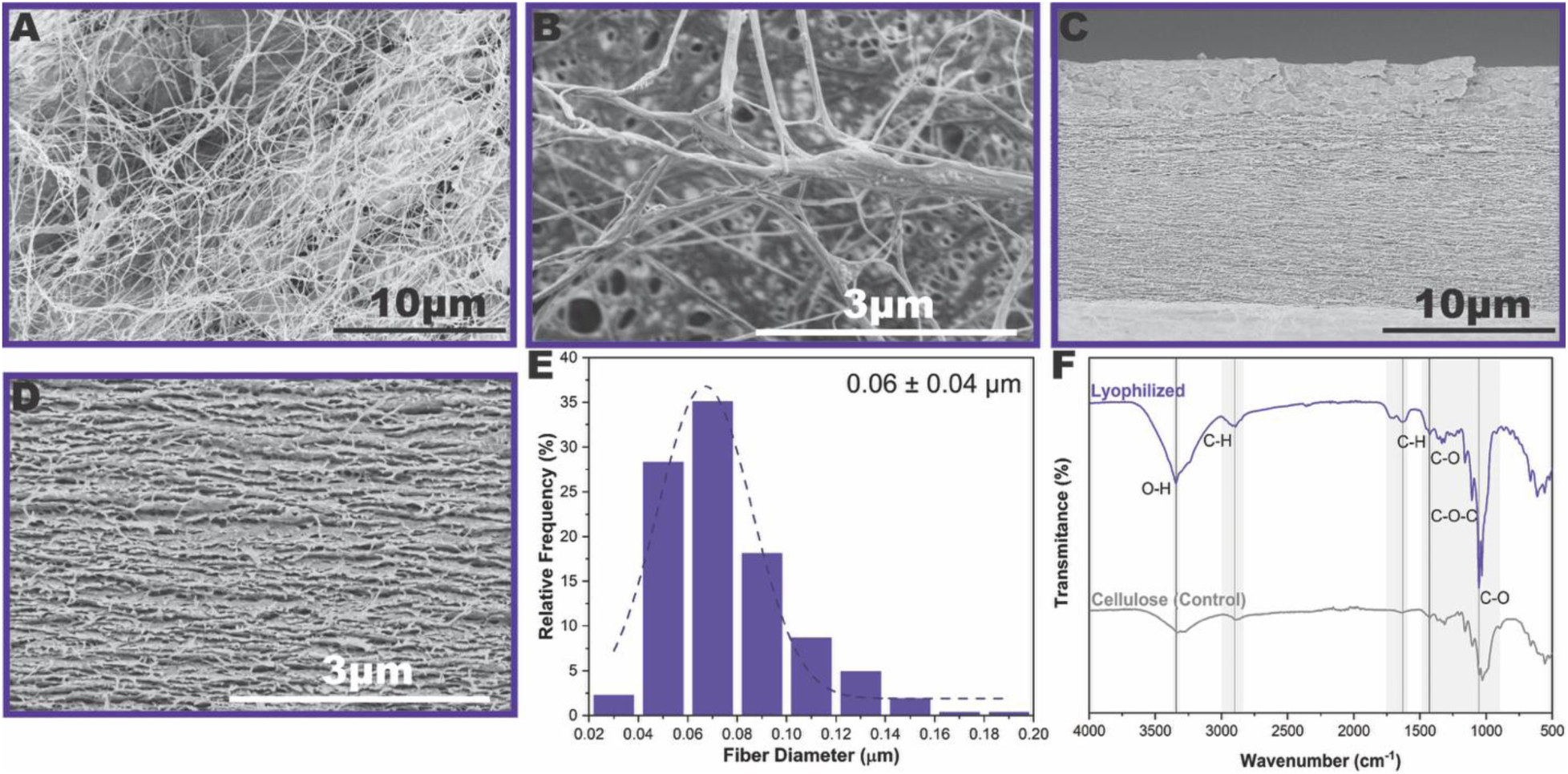
(**A-B**) SEM micrographs of kombucha-derived lyophilized non-wovens. **(C-D)** Cross-sectional SEM micrographs of lyophilized non-wovens. Scale bars **(A, C)** 10 μm and **(B, D)** 3 μm are provided. **(E)** Fiber diameter distribution fitted with a Gaussian curve. The average fiber diameter is provided in the upper right corner. **(F)** FTIR spectra of kombucha-derived lyophilized non-woven and cellulose powder (control).

**Figure 3F** presents a representative spectra of the lyophilized kombucha derived cellulose non-woven and cellulose powder (control).^50^ The lyophilized spectra exhibits a broad O-H stretching band around 3340 cm⁻¹ and strong C-H stretching peaks near 2890 cm⁻¹, reflecting the abundant hydroxyl and aliphatic groups of cellulose. Absorptions at 1160-1000 cm⁻¹ are observed, corresponding to C-O and C-O-C stretching modes in the glucopyranose rings. A sharp band near 897 cm⁻¹ is attributed to the β-1,4-glycosidic linkages of cellulose.^51^ A shoulder around 1645 cm⁻¹ is due to adsorbed water, and minor absorbance in the 1500-1700 cm⁻¹ region likely arises from trace phenolic carbonyls.^50^ The peaks at 1540-1550 cm⁻¹ and 1640-1650 cm⁻¹ (amide I and II) indicate the presence of microbial proteins. Overall, as expected, the kombucha derived non-wovens are primarily cellulose with some evidence of residual protein components.^52^

### Mechanical Performance of Different Non-Wovens Thicknesses

Next, we evaluated how fermentation time and thus, non-woven thickness influenced mechanical performance. We note that these studies unveiled an unexpected trend: although thicker non-wovens contained more accumulated cellulose, they did not exhibit increased mechanical properties, as will be described in detail in this section. Non-wovens were grown for either 7 days to an average thickness of 0.39 ±0.14 mm or for 21 days resulting in a thicker, 2.52 ±0.41 mm non-woven (**Table S2**). We named these samples “thin” and thick”. **Figure 4A** displays a representative plot acquired via tensile testing that displayed an initial linear-elastic region followed by yielding and eventual fracture. The thin non-wovens showed a steeper initial slope and a statistically higher stress than the thick non-wovens. Likewise, the elongation at break was higher for the thin non-wovens: thin non-wovens extended by 21.91 ±1.76% before failure, whereas thick non-wovens break at 25.23 ±3.30% strain (**Figure 4A-B**). The Young’s modulus for the thin non-wovens was 45.97 ±21.29 MPa, whereas the thick non-wovens have a statistically lower modulus of 7.323 ±4.454 MPa as shown on **Figure 4C**. In parallel with the modulus differences, the ultimate tensile strength was statistically greater for the thin than the thick non-wovens, 6.55 ±1.76 MPa versus 0.38 ±0.11 MPa, respectively. Interestingly, the thin non-wovens carry nearly an order of magnitude more stress before fracturing. Daus *et al.*^53^ likewise reported a thickness-dependent stiffness-ductility trade-off in purified non-wovens. Their thin mats, which were 1.4-1.7 mm in diameter exhibited a high elastic modulus of 2572 ±661 MPa but only a 1% strain at break, whereas their thicker 3-7 mm mats had a lower stiffness of 282 ±140 MPa paired with an 11% strain at break. While the values reported in that study were higher than those observed here, this might be attributed to post-processing, as will be discussed and benchmarked in later sections.

**Figure 4.**
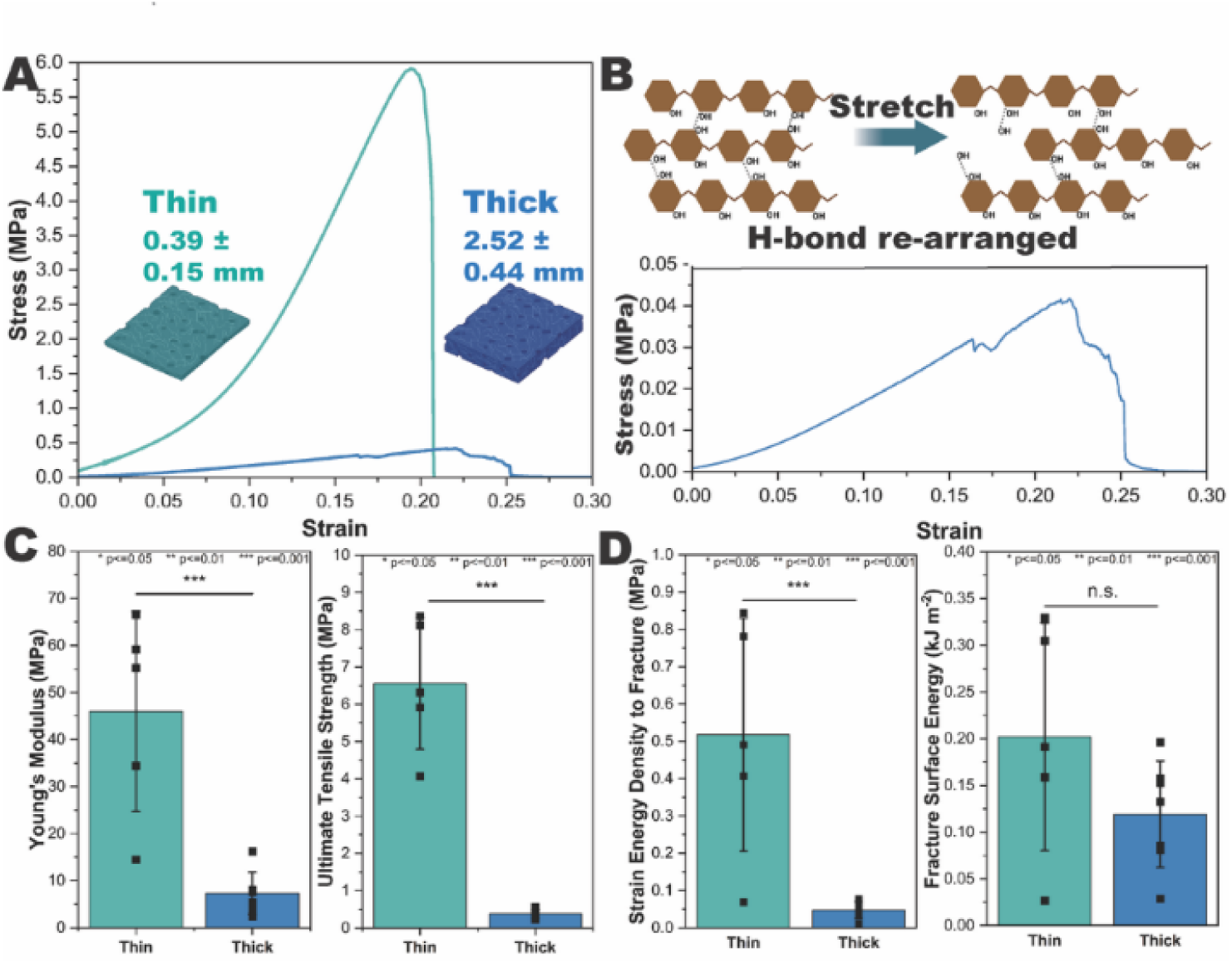
Representative stress-strain curves for **(A)** thin and thick non-wovens. **(B)** Schematic diagram of fiber re-arranged and magnified view of thick non-woven from figure **(A)**. **(C)** Young’s modulus and ultimate tensile strength, **(D)** strain energy density to fracture, and fracture surface energy for thin and thick non-wovens.

Next, we quantified the strain energy density to fracture and fracture surface energy as calculated using **Equations S1** and **S2**. In **Figure 4D**, the thin non-wovens exhibited a strain energy density to fracture of 0.52 ±0.3 MPa, whereas the thick showed a significantly lower value of 0.05 ±0.02 MPa, indicating reduced energy absorption per unit volume prior to failure. In contrast, the calculated fracture surface energies were 0.20 ±0.1 kJ m⁻² for thin and 0.12 ±0.1 kJ m⁻² for thick non-wovens, suggesting that the total energy required to generate new fracture surface was similar in magnitude. This difference likely arises because fracture surface energy scales with both strain energy density and sample thickness, such that the thicker non-wovens partially compensate for their lower energy density to fracture.

Cross-sectional SEM images in **Figure 3C-D** reveal a multilayered architecture that likely enhances energy absorption through interlayer sliding while simultaneously reducing interfacial cohesion and, in turn, stiffness.^54^ Moreover, prolonged fermentation may trap residual cells and organic byproducts within the network, disrupting hydrogen-bond percolation and plasticizing the hydrated matrix.^53^ Consistent with this interpretation, Daus *et al.*,^53^ reported that thicker kombucha-derived films retained more fermentation residues and water, leading to reduced stiffness. Thus, extended culture time introduces a trade-off between modulus and elongation.

### Mechanical Properties of Post-Growth Drying Treatment Non-Wovens

Next, how the mechanical behavior of wet kombucha-derived non-wovens was impacted by post-growth processing was explored. After oven-drying or lyophilization the non-wovens exhibited an equivalent decrease in thickness of >20% to ∼0.26 mm from ∼1.24, see data provided on **Table S2**. Uniaxial tensile tests shown on **Figure 5A** reveal that the wet non-wovens displayed an initial slope that began to yield almost immediately, displaying a long yielding or ductile region after a small elastic segment. The oven-dried curve lies in between: its initial slope is higher than the wet non-wovens but lower than that of the lyophilized non-wovens, and it shows a modest yield point. In contrast, the lyophilized non-wovens showed the steepest initial slope and remained nearly linear until fracture. In **Figure 5B**, the lyophilized non-wovens exhibited greater elongation at break (24.55 ±1.9%) than the wet which broke at 9.35 ±2.8% and oven-dried non-wovens which broke at 6.03 ±0.8%. The elastic regime extended to 1-2% strain for both dry conditions, with the lyophilized samples exhibiting a statistically significant increase (p<0.05). In contrast, the wet non-wovens departed from linearity at very low strain. This behavior reflects water-induced plasticization: absorbed water disrupts inter-fibrillar hydrogen bonding and reduces frictional resistance between fibers, thereby lowering stiffness and promoting early yielding.^55^

**Figure 5.**
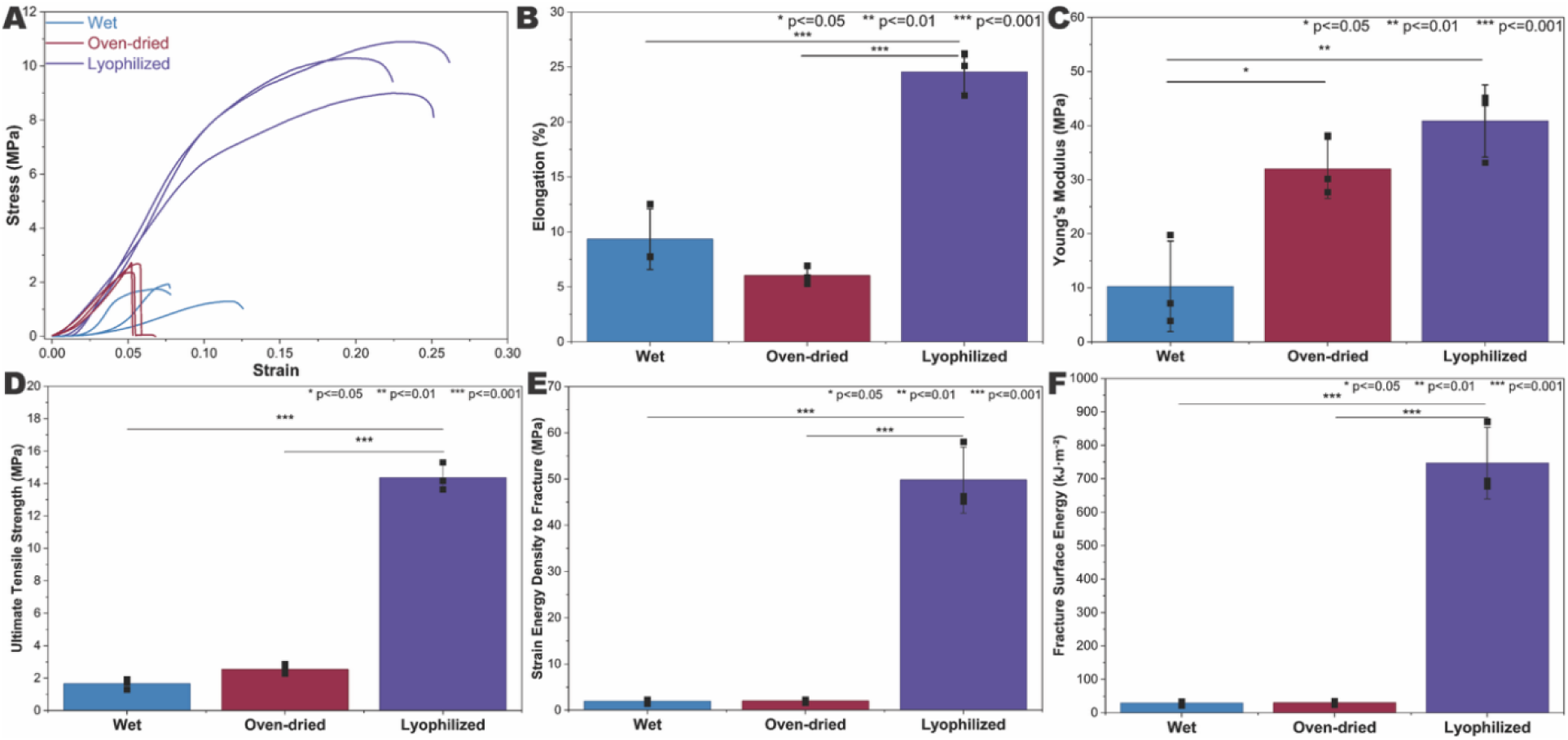
Plots displaying the **(A)** stress-strain, **(B)** elongation, **(C)** Young’s modulus, **(D)** ultimate tensile strength, **(E)** strain energy density to fracture, and **(F)** fracture surface energy of wet, oven-dried, and lyophilized kombucha derived non-wovens.

**Figure 5C** compares the average Young’s modulus of the non-wovens. The lyophilized non-wovens displayed the highest modulus (40.84 ±6.7 MPa), which was statistically greater than the wet (10.27 ±8.37 MPa) and oven-dried (31.99 ±5.5 MPa). Post-treatment produces a higher modulus material, whereas the presence of moisture reduced the stiffness. Zeng *et al.*^56^ reported that dried bacterial cellulose films exhibited significantly higher Young’s modulus and tensile strength than freeze-dried films. Conversely, water has a pronounced plasticizing effect on cellulose; fully hydrated bacterial cellulose networks displayed a dramatically lower stiffness due to disrupted hydrogen bonding, as reported by Rebelo *et al*.^57^ **Figure 5D** shows that the wet non-wovens had an ultimate tensile strength of 1.66 ±0.3 MPa, which was statistically equivalent (p>0.05) to the oven-dried at 2.54 ±0.29 MPa. In contrast, the lyophilized non-wovens had a statistically greater ultimate tensile strength which was 14.36 ±0.9 MPa.

**Figure 5E** presents the strain energy density to fracture for each condition. The wet and oven-dried non-wovens displayed an equivalently low toughness ∼2 MPa. The lyophilized non-wovens absorb statistically more energy (49.80 ±7.2 MPa. In practical terms, the lyophilized non-woven withstand much more mechanical work before failing. **Figure 5F** shows the fracture surface energy for the three conditions. The lyophilized non-wovens exhibited a strain energy density of 746.97 ±107.2 kJ/m^2^, which was statistically greater than the wet (291.55 ±686.5 kJ/m^2^) or oven-dried (298.53 ±486.9 kJ/m^2^) non-wovens. Like the work-of-fracture, the lyophilized value is significantly larger than the others (p<0.001), whereas wet and oven-dried mats are equivalent (no significant difference). This confirms that the lyophilized mats fail in a more energy-dissipating (tough) manner, whereas the other samples exhibit brittle failure with minimal energy absorption.

We were also curious to explore the impact of re-hydrating the oven-dried and lyophilized non-wovens on their mechanical performance. **Table S2** shows that upon re-exposure to water, both the oven-dried and lyophilized samples re-gained 84% and 85% of their initial wet thickness. **Figure S1** shows that upon re-hydration, the Young’s modulus of the oven-dried mats stayed statistically equivalent, whereas their ultimate tensile strength greatly increased to 6.947 ±0.35 MPa. Interestingly, re-hydrating the lyophilized non-woven led to a significant improvement in both their Young’s modulus and ultimate tensile strength (9.584 ±0.75 MPa). Overall, this suggests that that the non-wovens can be stored in a dry state and upon rehydration, provide excellent mechanical properties.

To further probe network stability under deformation, we employed rheology to characterize small-strain viscoelastic behavior, complementing the large-strain response measured by tensile testing. Frequency sweep tests (**Figure 6A**) reveal a predominantly elastic behavior (G′ ≫ G″) across 0.1-100 rad s**^-1^** for all kombucha-derived non-wovens, but with different stiffness depending on the drying process. The lyophilized non-wovens consistently show the highest G’ (3.4×10^5^ Pa), which was an order of magnitude greater than the wet (1.5×10⁵ Pa) and oven-dried (4.6×10⁴ Pa) non-wovens, data provided in **Table S3**. Oscillation amplitude strain sweep displayed on **Figure 6B** identified the linear viscoelastic region, defined as the strain at which G’ deviated by ≤5% from its plateau. Lyophilized non-wovens showed the broadest linear viscoelastic region, with a nearly constant G′ up to 0.4 strain and the highest plateau G′ (1.1×10^6^ Pa). Wet non-wovens reached the linear viscoelastic region limit at 0.05 strain but showed a more gradual modulus decay thereafter, consistent with lubricated fibril sliding. Oven-dried non-wovens remained linear up to 0.2 strain followed by modulus loss consistent with stress softening previously reported in hydrogen-bonded cellulose networks.

**Figure 6.**
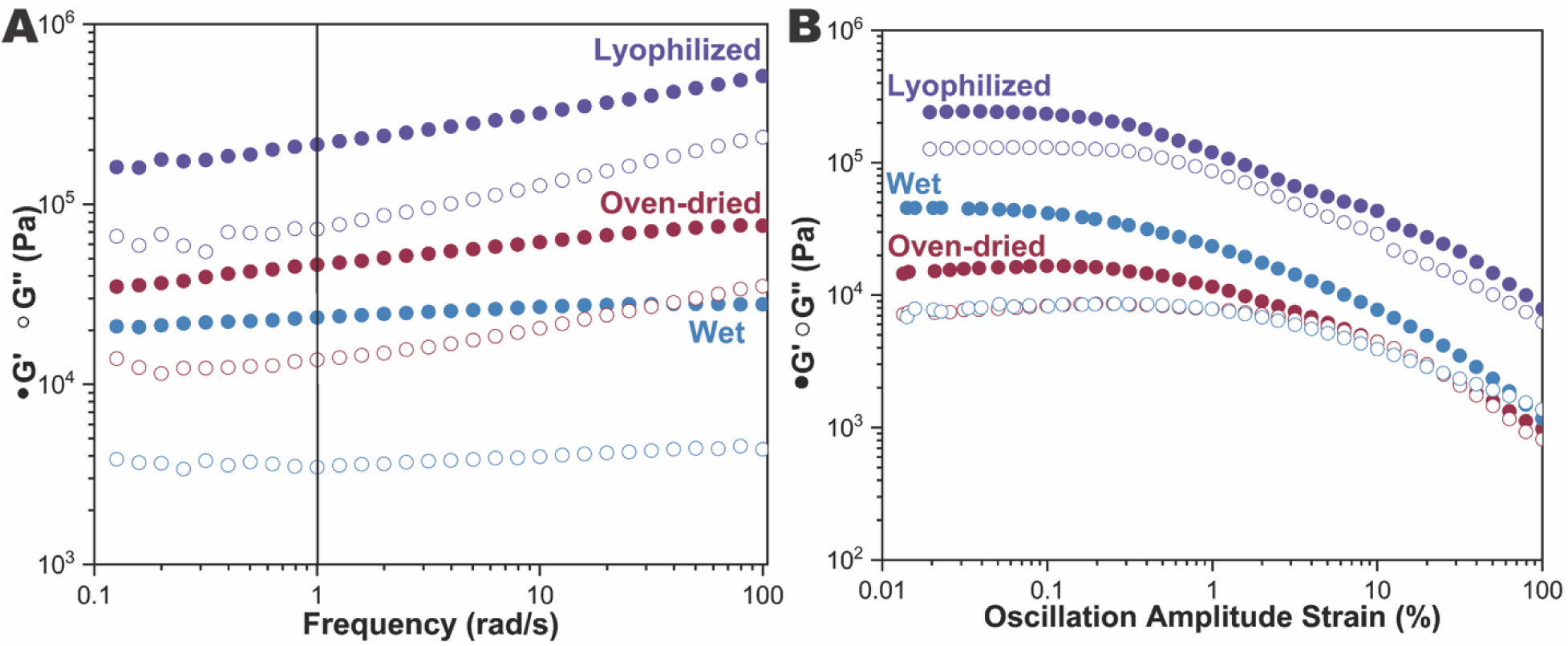
Rheological characterization of kombucha derived non-wovens (wet, lyophilized and oven-dried) including: **(A)** frequency sweep showing G′ and G″ over a range of angular frequencies and **(B)** oscillation amplitude strain sweep showing G′ and G″ as a function of strain.

### Discussion on the Mechanical Properties of Kombucha Derived Non-Wovens

**Figures 5** and **6** show that processing alters the mechanical properties of kombucha derived non-wovens. To summarize the literature, in **Figure 7**, we have plotted the ultimate tensile strength and elastic modulus of our non-wovens, as well as literature-reported cellulose non-wovens. More details on what has been plotted are provided on **Table S4.** All hydrated cellulose materials are clustered in the low-stiffness, low-strength regime, as indicated by a blue shaded area on **Figure 7**. The wet non-wovens are soft and compliant due to the water present, which promotes fiber sliding. Our lyophilized non-woven showed a ∼4× higher Young’s modulus and a ∼9× higher tensile strength than they were as-prepared (wet). Lyophilization differs in that water is removed by sublimation, which helps preserve the porous nanofiber network and avoid capillary-driven collapse, yielding a material that is both stiff and relatively extensible.^45^ Fatima *et al.* reported a tensile strength of 9.2 ±3.5 MPa and an elastic modulus of 68.6 ±5.2 MPa for a lyophilized bacterial cellulose membrane produced by *Komagataeibacter sp*. In comparison, oven-dried samples underwent evaporative drying, where capillary forces can densify or partially collapse the fiber network. Consistent with the report from Andree *et al*.^58^ oven drying results in a higher modulus mat at the expense of elongation at break. These values further support the trend that sublimation-based drying can substantially increase stiffness and strength relative to hydrated cellulose materials.

**Figure 7.**
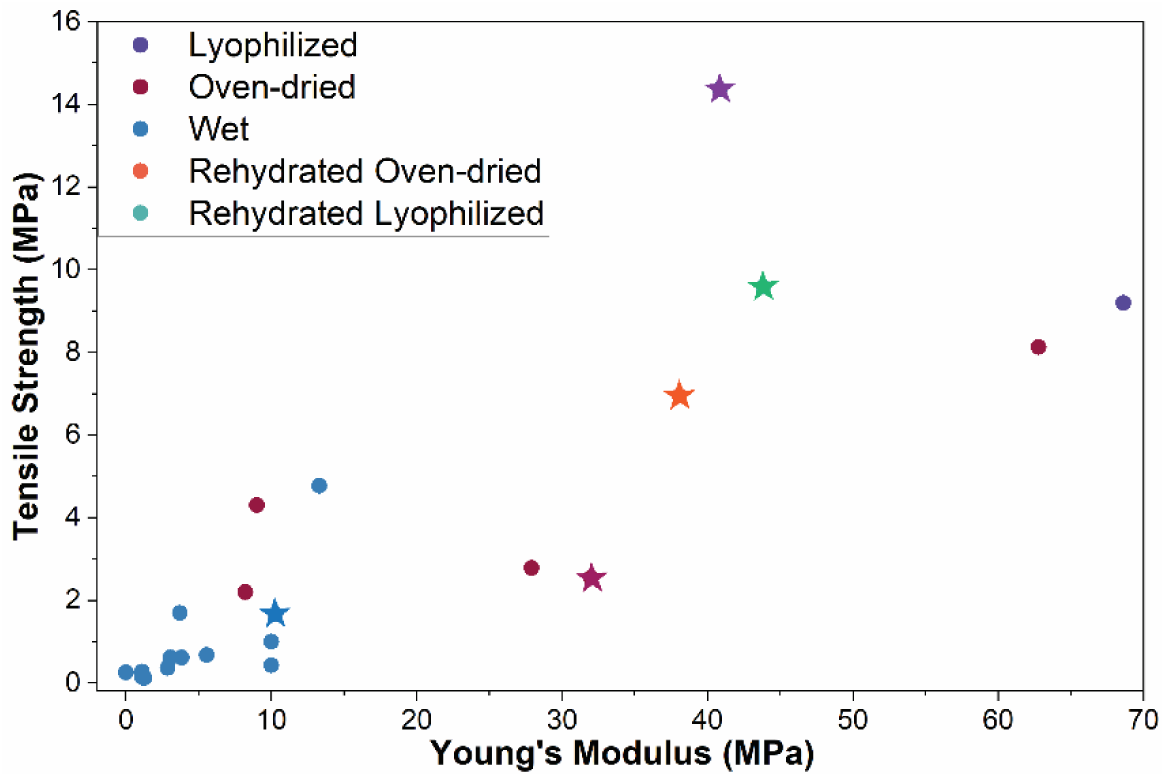
Summary of literature reported mechanical properties of cellulose non-wovens. The triangles represent data from this work. The full dataset is provided in **Table S4**.

### Proof-of-Concept Recycling of Kombucha Derived Non-Wovens

As a proof-of-concept demonstration of circularity, we investigated whether our kombucha-derived cellulose could be reprocessed through a closed-loop materials cycle. As illustrated in **Figure 8A**, kombucha-derived non-wovens were first fabricated into infant onesies that, once outgrown, were enzymatically degraded within 72 hr (**Figure 8B**). **Figure S2** and **Table S5** provides the FTIR spectra and peak assignments that support that the kombucha derived non-wovens underwent partial hydrolysis. The degradation product retained strong carbohydrate bands in the 1150-1000 cm⁻¹ region, consistent with glucose-rich species, while the cellulose-associated bands near ∼1430 cm⁻¹ and ∼897 cm⁻¹ were not obvious, consistent with partial disruption of the cellulose chains. The resulting products were then blended with an aqueous PVA solution and electrospun to form a second-generation fibrous mat (**Figures 8C** and **S3**). During spinning, calcofluor white stain was introduced, which is a fluorescent dye that binds specifically to β-1,4 and β-1,3 glycosidic linkages. The continuous blue fluorescence that we observed along the fibers strongly supports the presence of β-glycosidic linkages and other cellulose-oligosaccharides present throughout the fibrous network.

**Figure 8.**
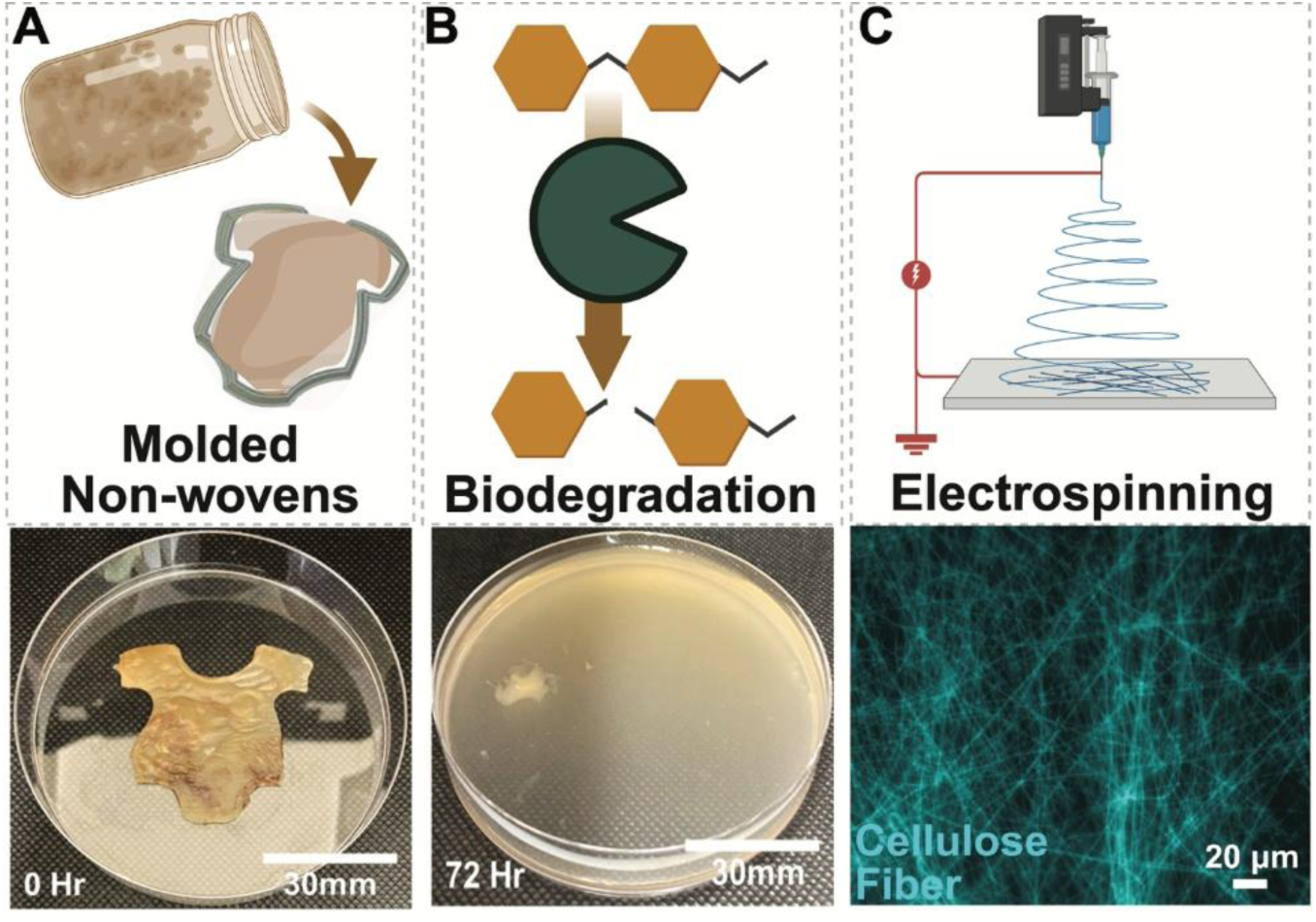
Top row includes schematic representations, while the bottom row provides the images of **(A)** non-wovens grown from kombucha, **(B)** cellulase-mediated hydrolysis of cellulose into glucose units after 72 hr, and **(C)** the re-manufacturing via electrospinning of the degradation products into new fibrous mats. Calcofluor white stain was used and a scale bar of 50 μm is provided.

This proof-of-concept recycling experiment is included as a demonstration of the reprocessing potential of robust kombucha-derived textiles into other materials. The scale of textile waste and petroleum use motivates the search for alternative chemistries, fabrics, and green manufacturing processes that decreases our environmental impact. We suggest that kombucha-derived bacterial cellulose offers a sustainable advantage over petroleum-based textiles. These results establish a closed-loop pathway in which kombucha-derived cellulose non-wovens can be grown, used, enzymatically degraded, and reassembled into a new fibrous material.

## CONCLUSION

In this study, we explored the connection between kombucha-derived cellulose non-woven fermentation conditions, non-woven post-processing, mechanical performance, and circular reusability. By systematically varying inoculum concentration, sucrose concentration, tea content, temperature, and initial pH value, we demonstrate that non-woven thickness can be tuned through fermentation conditions. Variations in thickness directly influenced mechanical behavior; the thin non-wovens were stronger, with a Young’s modulus of 4.60 ±2.1 MPa and an elongation at break of 21.91 ±1.8%, versus the thick with a modulus of 0.73 ±0.5 MPa and elongation of 25.23 ±3.3%. The mechanical properties were further shaped by processing history, such as lyophilization which demonstrated the highest modulus of 40.84 ±6.7 MPa. Finally, we demonstrated a closed-loop reuse strategy in which kombucha-derived non-wovens were enzymatically depolymerized into glucose-rich hydrolysates and cellulose fragments, followed by re-manufacturing through electrospinning into second-generation non-woven scaffold. This paper demonstrates the potential of using bio-manufacturing (*i.e.,* fermentation) and the bio-based feedstock of cellulose as a circular path forward towards the fermentation of new materials with controllable materials properties. We suggest that kombucha-derived bacterial cellulose offers a sustainable advantage over petroleum-based textiles for use in wearable devices and clothing, separation membranes, green packaging, and wound dressings.

## ASSOCIATED CONTENT

### Supporting Information

Experimental procedures; fermentation studies, thickness measurements; tensile testing and rheological data; benchmarking data; FTIR spectra and peak assignment tables.

## AUTHOR INFORMATION

### Corresponding Author

* Jessica D. Schiffman, Department of Chemical and Biomolecular Engineering, Materials Science and Engineering Graduate Program, University of Massachusetts Amherst, Amherst, Massachusetts 01003-9303, USA.

### Author Contributions

The manuscript was written through contributions of all authors. All authors have given approval to the final version of the manuscript.

### Notes

The authors declare no competing financial interest.

## ACKNOWLEDGMENT

This work was supported in part by a Fellowship from the University of Massachusetts Amherst to L.H. through the Biotechnology Training Program (National Research Service Award T32 GM135096). L.H. also acknowledges support from the Materials Science and Engineering Doctoral Fellowship. We acknowledge the support of the National Science Foundation (Award #2150075). We thank Dr. Shao-Hsiang Hung and B.F. Coleman for SEM assistance at the W.M. Keck Center for Electron Microscopy. Thanks to Dr. Alfred Crosby (UMass Amherst) for access to their texture analyzer and CO₂ laser cutter.

## ABBREVIATIONS

ANOVA: analysis of variance
FTIR: Fourier-transform infrared spectroscopy
CFU: colony-forming unit
CFW: calcofluor white
CesA: cellulose synthase A
CO₂: carbon dioxide
DI: deionized
PBS: phosphate-buffered saline
PES: polyethersulfone
PVA: poly(vinyl alcohol)
SAOS: small-amplitude oscillatory shear
SCOBY: symbiotic culture of bacteria and yeast
SD: standard deviation
SEM: scanning electron microscopy
UTS: ultimate tensile strength
YPD: yeast extract-peptone-dextrose

## FOR TABLE OF CONTENTS ONLY

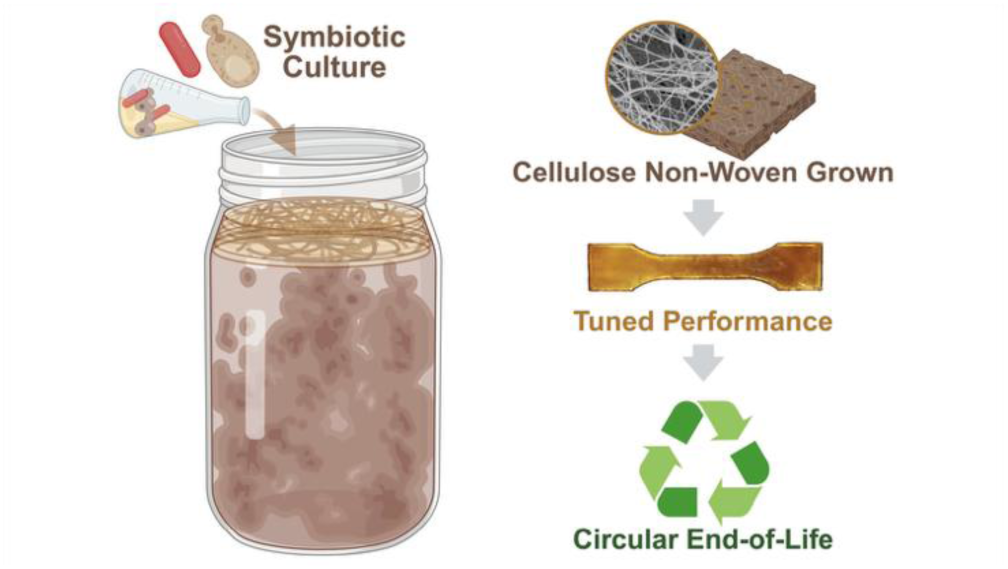

## Notes

### Competing Interest Statement

The authors have declared no competing interest.

