## Supplementary material for "Kombucha-Derived Cellulose Non-wovens: Growth Optimization, Mechanics, and Recycling": SI file

**Supplementary Methods: Mechanical Testing of Kombucha-Derived Non-Wovens.**

**Oscillatory Rheological Method and Data Analysis**. The amplitude (strain) sweep identified the linear viscoelastic region (LVR). The reported LVR limit corresponds to the strain at which G′ deviated by ≤5% from its plateau. Frequency sweeps were then run within the LVR (0.1 to 100 rad $s^{-1}$ at a fixed strain ≤ the LVR limit for each state). G′ at 1 rad $s^{-1}$ is reported from the frequency sweep. Values summarize independent specimens per state.

**Uniaxial Tensile Method and Data Analysis.** Young's modulus, $E$, was determined by linear regression of the initial linear regime of the engineering stress-strain response. Ultimate tensile strength (UTS), $\sigma_{max}$, was defined as the maximum engineering stress reached prior to failure, and elongation at break, $\varepsilon_b$, was taken as the engineering strain at failure. The strain energy density to fracture, $W$, was defined as the area under the engineering stress-strain curve up to the point of failure as **Equation S1**:

$$W = \int_0^{\varepsilon_b} \sigma(\varepsilon)\, d\varepsilon \quad \textbf{(S1)}$$

where $W$ is the strain energy density to fracture, $\sigma(\varepsilon)$ is the engineering stress at strain $\varepsilon$, $\varepsilon$ is the engineering strain, and $\varepsilon_b$ is the strain at break. Numerical integration was performed using the trapezoidal rule. The fracture surface energy, $G$, defined as the energy dissipated per unit newly created fracture surface area, was estimated using **Equation S2**:

$$G = tW \quad \textbf{(S2)}$$

where $G$ is the fracture surface energy, $t$ is the sample thickness, and $W$ is the strain energy density to fracture. This approach assumes uniform energy dissipation across the fracture plane and follows common practice for estimating fracture energies of soft and fibrous materials from tensile measurements.

**Table S1.** Influence of Culture Variables on the Thickness of Kombucha-Derived Non-Wovens.

| **Culture variables** | | **Thickness** (mm) | **Normalized Ratio†** |
|---|---|---|---|
| **Inoculation** ($L \cdot L^{-1}$) | 0.01 | 0.096 ±0.1 | 0.00 |
| | 0.02 | 0.064 ±0.04 | -0.33 |
| | 0.05 | 0.36 ±0.06 | 2.75 |
| | 0.1 | 1.769 ±0.1 | 17.43 |
| **Sucrose** ($g \cdot L^{-1}$) | 0 | 0.003 ±0.001 | 0.00 |
| | 50 | 0.76 ±0.2 | 252.33 |
| | 100 | 2.08 ±0.2 | 692.33 |
| | 150 | 2.34 ±0.3 | 779.00 |
| **Tea leaves** ($g \cdot L^{-1}$) | 5 | 0.36 ±0.1 | 0.00 |
| | 10 | 0.69 ±0.04 | 0.92 |
| | 15 | 1.11 ±0.5 | 2.08 |
| | 20 | 0.95 ±0.1 | 1.64 |
| **Temperature** (°C) | 20 | 1.25 ±0.1 | 0.00 |
| | 25 | 1.07 ±0.05 | -0.14 |
| | 30 | 1.68 ±0.2 | 0.34 |
| | 35 | 0.13 ±0.06 | -0.90 |
| **pH Value** | 3 | 0.20 ±0.07 | 0.00 |
| | 4 | 1.35 ± 0.1 | 5.75 |
| | 5 | 0.08 ±0.02 | -0.60 |
| | 6 | 0.01 ±0.0 | -0.95 |

†Calculated using **Equation 2**

**Table S2.** Thickness of Non-Wovens Under Different Fermentation, Drying, and Rehydration Conditions.

| **Sample Name** | **Fermentation Time** (days) | **Thickness** (mm) | **Normalized Ratio*** |
|---|---|---|---|
| Wet | 14 | 1.24 ±0.1 | 0.00 |
| Thin | 7 | 0.39 ±0.1 | -68.55 |
| Thick | 21 | 2.52 ±0.4 | 103.23 |
| Oven-dried | 14 | 0.26 ±0.1 | -79.03 |
| Lyophilized | 14 | 0.27 ±0.04 | -78.23 |
| Rehydrated Oven-dried | 14 | 1.04 ±0.1 | -16.13 |
| Rehydrated Lyophilized | 14 | 1.05 ±0.08 | -15.32 |

*As calculated using **Equation 2**

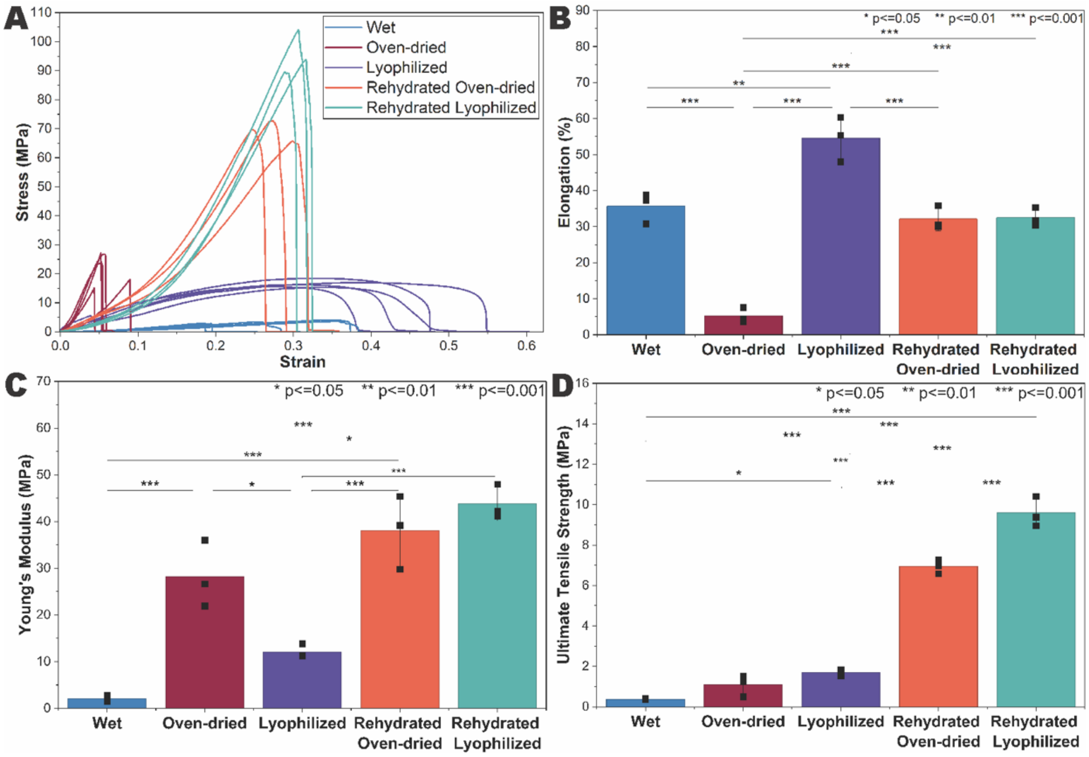


**Figure S1. (A)** Stress-strain curves, **(B)** elongation, **(C)** Young's modulus, and **(D)** ultimate tensile strength of various kombucha-derived non-wovens. Error bars denote one standard deviation (n = 3).

**Table S3.** Viscoelastic Response of Kombucha-Derived Non-Wovens.

| **Sample Name** | **G′ at 1 rad s$^{-1}$** | **LVR Limit in Strain Sweep (%)** |
|---|---|---|
| Wet | $1.5 \times 10^5$ | 0.05 |
| Oven-dried | $4.6 \times 10^4$ | 0.2 |
| Lyophilized | $3.4 \times 10^5$ | 0.4 |

**Table S4.** Benchmarking Mechanical Properties of Cellulose Non-Wovens.

| Sample Details | | Tensile Strength (MPa) | Young's Modulus (MPa) | Ref |
|---|---|---|---|---|
| Lyophilized | **Kombucha-derived non-wovens** | **14.36 ±0.9** | **40.84 ±6.7** | **This work** |
| | Bacterial cellulose membrane (*Komagataeibacter sp.*) | 9.2 ±3.5 | 68.6 ±5.2 | [1] |
| Oven-dried | **Kombucha-derived non-wovens** | **2.54 ±0.3** | **31.99 ±5.5** | **This work** |
| | Unmodified cellulose non-woven | 4.3 ±0.5 | 9.0 ±1.6 | [2] |
| | Wet unmodified cellulose non-woven | 2.2 ±0.1 | 8.2 ±0.4 | [2] |
| | Bacterial cellulose biofilm | 2.78 | 27.91 | [3] |
| | Bacterial cellulose membrane | 8.126 | 62.776 | [4] |
| | Crude dry bacterial cellulose | 11.6 ±0.8 | 180.3 ±10.6 | [5] |
| Wet | **Kombucha-derived non-wovens** | **1.66 ±0.3** | **10.27 ±8.4** | **This work** |
| | Bacterial cellulose hydrogel (*G. hansenii* CGMCC 3917) | 0.26 | 0.005 | [6] |
| | Bacterial cellulose membrane (*Komagataeibacter sp.* ATCC 700178) | 0.15 ±0.08 | 1.10 ±0.4 | [7] |
| | Bacterial cellulose membrane (*Komagataeibacter sp.* ATCC 10245) | 0.36 ±0.08 | 2.87 ±1.3 | [7] |
| | Bacterial cellulose membrane (*Komagataeibacter sp.* ATCC 23769) | 0.12 ±0.04 | 1.26 ±0.7 | [7] |
| | Bacterial cellulose membrane (*Komagataeibacter sp.* NBRC 13693) | 0.62 ±0.2 | 3.08 ±0.7 | [7] |
| | Bacterial cellulose membrane (*Komagataeibacter sp.* ATCC 53524) | 0.68 ±0.1 | 5.56 ±2.3 | [7] |
| | Bacterial cellulose membrane (*Komagataeibacter sp.* KTH 5655) | 0.62 ±0.2 | 3.83 ±1.1 | [7] |
| | Bacterial cellulose membrane | 4.770 | 13.305 | [4] |
| | Bacterial cellulose membrane (pH 5.0 Glycerol culture) | 1.70 | 3.70 | [7] |
| | Bacterial cellulose membrane (Glucose culture) | 0.12 | 1.26 | [6] |
| | Bacterial cellulose hydrogel | 0.12-0.68 | 1.10-5.56 | [8] |
| | Bi-networked cellulose nanohydrogel (clicked chemical cross-linking) | 0.43 | 10 | [9] |
| | Bacterial cellulose membrane | 1 ±0.0 | 10 ±0.0 | [8] |
| | Kombucha SCOBY | 0.27 ±0.08 | 1.09 ±0.4 | [11] |

| | | | | |
|---|---|---|---|---|
| | Crude fresh bacterial cellulose | 1.2 ±0.2 | 3.1 ±0.5 | [5] |
| | non-dried bacterial cellulose | 1.18 ± 1.01 | | |
| Rehydrated Oven-dried | **Kombucha-derived non-wovens** | **6.95 ±0.3** | **38.09 ±7.9** | **This work** |
| | Crude bacterial cellulose | 127.2 ±13.9 | 25.1 ±1.8 | [5] |
| | 25 °C convection-dried, then soaked | 17.40 ± 3.68 | | [12] |
| | 40 °C convection-dried, then soaked | 16.98 ± 4.34 | | [12] |
| | 80 °C convection-dried, then soaked | 8.49 ± 5.47 | | [12] |
| | 105 °C convection-dried, then soaked | 5.23 ± 1.59 | | [12] |
| Rehydrated Lyophilized | **Kombucha-derived non-wovens** | **9.58 ±0.7** | **43.83 ±3.7** | **This work** |
| | Freeze-dried, then soaked | 10.04 ± 2.32 | | [12] |

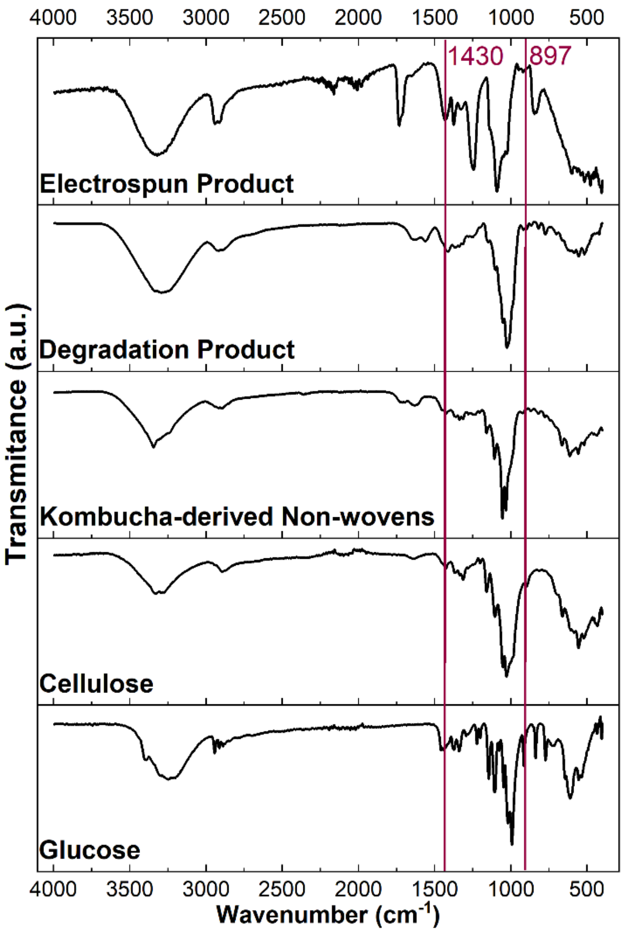


**Figure S2.** FTIR spectra of glucose, cellulose, kombucha-derived cellulose non-wovens, enzymatic degradation product, and electrospun fibers. The peaks at ~1430 $cm^{-1}$ and ~897 $cm^{-1}$ indicate cellulose-associated bands assigned to $CH_2$ bending and the β-(1→4)-glycosidic linkage, respectively.

**Table S5.** FTIR Peak Assignments for Cellulose, Glucose, and Kombucha-Derived Samples.

| **Wavenumber** ($cm^{-1}$) | **Assignment** | **Glucose** | **Cellulose** | **Kombucha Non-wovens** | **Degradation Product** | **Electrospun Product** |
|---|---|---|---|---|---|---|
| ~3330 | O-H stretching[13] | Yes | Yes | Yes | Yes | Yes |
| ~2900 | C-H stretching[14] | Yes | Yes | Yes | Yes | Yes |
| ~1430 | $CH_2$ bending[14] | No | Yes | Yes | Yes | Yes |
| ~897 | β-(1→4) glycosidic vibration[14] | No | Yes | Yes | Yes | Yes |
| 1655 / 1540 | Amide I / II[15] | No | No | Yes | No | No |
| 1150-1000 | C-O / C-C stretching[14] | Yes | Yes | Yes | Yes | Yes |

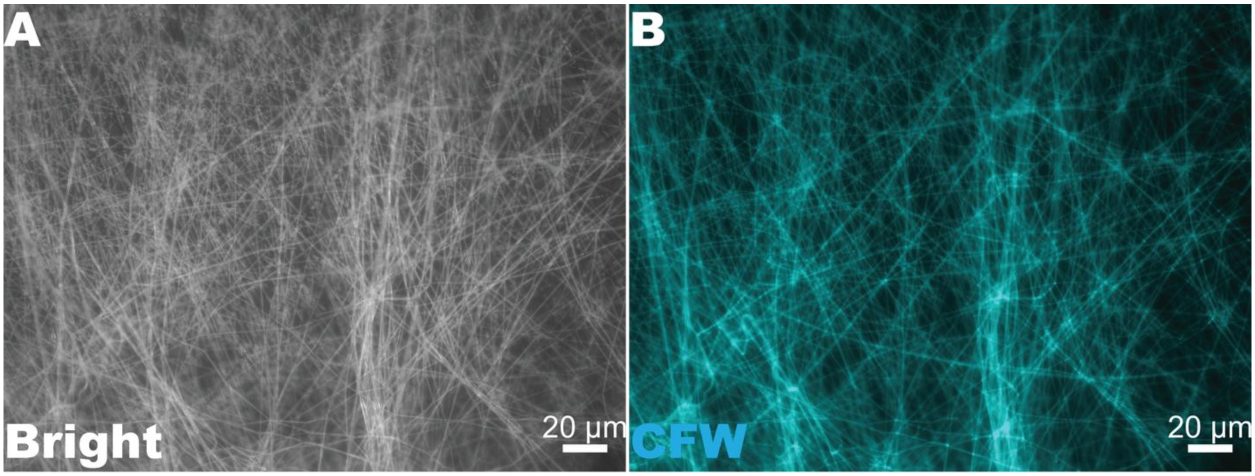


**Figure S3**. **(A)** Brightfield and **(B)** fluorescence images of electrospun fibers prepared from kombucha-derived hydrolysis products. Scale bars: 20 µm.
